# Dynamic regulatory phosphorylation of mouse CDK2 occurs during meiotic prophase I

**DOI:** 10.1101/2023.07.24.550435

**Authors:** Rachel A. Bradley, Ian D. Wolff, Paula E. Cohen, Stephen Gray

## Abstract

During prophase I of meiosis, DNA double-strand breaks form throughout the genome, with a subset repairing as crossover events, enabling the accurate segregation of homologous chromosomes during the first meiotic division. The mechanism by which DSBs become selected to repair as crossovers is unknown, although the crossover positioning and levels in each cell indicate it is a highly regulated process. One of the proteins that localises to crossover sites is the serine/threonine cyclin-dependent kinase CDK2. Regulation of CDK2 occurs via phosphorylation at tyrosine 15 (Y15) and threonine 160 (T160) inhibiting and activating the kinase, respectively. In this study we use a combination of immunofluorescence staining on spread spermatocytes and fixed testis sections, and STA-PUT gravitational sedimentation to isolate cells at different developmental stages to further investigate the temporal phospho regulation of CDK2 during prophase I. Western blotting reveals differential levels of the two CDK2 isoforms (CDK2^33kDa^ and CDK2^39kDa^) throughout prophase I, with inhibitory phosphorylation of CDK2 at Y15 occurring early in prophase I, localising to telomeres and diminishing as cells enter pachynema. Conversely, the activatory phosphorylation on T160 occurs later, specifically the CDK2^33kDa^ isoform, and T160 signal is detected in spermatogonia and pachytene spermatocytes, where it co-localises with the Class I crossover protein MLH3. Taken together, our data reveals intricate control of CDK2 both with regards to levels of the two CDK2 isoforms, and differential regulation via inhibitory and activatory phosphorylation.

## Introduction

During meiosis, DNA must be accurately segregated via two rounds of cell division, first in a reductional division segregating homologous chromosomes, and then as an equational division segregating sister chromatids. If successful, meiosis leads to the production of viable haploid gametes (sperm and egg). Crossover events, the exchange of genetic material between parental homologous chromosomes, are required to physically link each chromosome pair enabling their accurate segregation at the first meiotic division. Crossovers are generated through meiotic recombination, where DNA double-strand breaks (DSBs) are generated throughout the genome through the action of the topoisomerase-like protein SPO11, its partner TOPOVIBL, and multiple essential accessory proteins during early leptonema^1-3^. Following removal of SPO11 and resection of the DNA ends^4,5^, DSBs can enter different repair pathways, with the recruitment and activity of specific DNA repair proteins defining the potential repair outcomes (reviewed in ^6^).

In the mouse, 200-300 DSBs form in each cell, with most (approximately 90%) repairing as non-crossover events, and the remaining 23-26 DSB events repairing as crossovers. Of the crossovers that form, the vast majority occur through the Class I/ZMM pathway, resolved by the MutLγ endonuclease complex consisting of MutL homologs, MLH1 and MLH3^7-10^. The remaining crossovers form through the Class II, MUS81-EME1 dependent pathway and a yet to be determined additional pathway(s)^11^. The genome-wide distribution of crossovers must be exquisitely controlled to ensure that every homologous chromosome pair receives at least one crossover (termed the obligate crossover)^12^, that crossovers that form do not occur close to each other (crossover interference, only affecting Class I crossovers)^13-15^, and that crossovers are maintained at the expense of non-crossovers when DSBs are limiting (crossover homeostasis)^16,17^. However, the exact mechanism that defines which DSBs are selected to become crossovers is not well understood.

Characterisation and analysis of meiotic mutants and protein localisation by immunofluorescence labelling on spread spermatocytes has revealed the strict temporal dynamics of key pro-crossover factors. Of the DSBs that form, half (approximately 150) recruit the mismatch repair family proteins MSH4/MSH5 (MutSγ complex) in zygonema^18,19^, with the number of MutSγ foci decreasing to approximately 50-60 as cells reach mid-pachynema. Regulation of MutSγ foci is dependent upon the SUMO E3 ligase RNF212, with less than half of the final MutSγ foci co-localising with RNF212^20^. By pachynema, additional proteins co-localise within this complex, including the E3 Ubiquitin ligase HEI10^21,22^, cyclin-domain containing protein CNTD1^23^, the CNTD1 interacting protein PRR19^24^, the aforementioned MutLγ endonuclease complex^7,9,10,19^, and the cyclin-dependent kinase CDK2^25^.

CDK2 is a serine/threonine kinase that interacts with A and E type cyclins^26^. Given the localisation of cyclin-domain containing protein CNTD1 and CDK2 at crossover sites, it was proposed that CNTD1 may function as a meiosis specific cyclin. The CNTD1 ortholog in *C. elegans*, COSA-1, mediates crossover formation in worm oocytes by promoting CDK-2 phosphorylation of MSH-5^27^, suggesting a conserved role for CNTD1/COSA-1 in regulating CDK-2 activity. However, *in vivo* data suggest this is not the case in mouse spermatocytes since CNTD1 fails to interact with CDK2^23,24^, and therefore it is unclear how CDK2 accumulation to crossover sites is controlled. In addition to localisation at crossover sites during pachynema, CDK2 localises to telomeres from zygonema through to diplonema. The telomeric interaction with the Speedy/RINGO family protein Speedy A modulates interactions with the telomere and nuclear envelope^28-30^. CDK2 protein, and its kinase activity, is critical for germ cell development, with *Cdk2*^*-/-*^ mice arresting in early prophase I^31-34^.

CDK2 kinase activity is regulated not only by its interaction with Cyclins, but also by phosphorylation. The WEE1 kinase phosphorylates CDK2 on threonine 14 (T14) and tyrosine 15 (Y15), inhibiting its activity^35^. Counter to WEE1 action, phosphorylation by <u>C</u>dk-<u>a</u>ctivating <u>k</u>inase complex (CAK) CDK7-Cyclin H on threonine 160 (T160) activates CDK2^35,36^. In addition, the phosphatase CDC25 also regulates CDK2 by removing inhibitory phosphorylation events, enabling its activation^37-39^. Because *Cdk2*^*-/-*^ mice arrested in early prophase I before pachynema, the role of CDK2 during crossover formation remained elusive. However, recent work using mice containing a separation of function mutation of the activatory T160 residue on CDK2 (T160A) showed normal telomeric association of CDK2 and proper homolog synapsis but failed to load Class I crossover marker MLH1, indicating that activated CDK2 is essential for crossover formation^40^. In addition, a “hyperactive” form of CDK2 generated through a mutation of the inhibitory Y15 site (Y15S) resulted in elevated interstitial CDK2 and MLH1 foci on autosomes^40^. To further characterise the regulation of CDK2 in mouse spermatocytes, we focused our investigation on these two important residues.

Interestingly, CDK2 exists in two isoforms in mouse testis, a 33kDa and a 39kDa form^41^, and the relative contribution of each of these isoforms to CDK2’s role during prophase I to promote homolog synapsis and crossover formation has not been investigated. In addition, the temporal regulation and spatial localisation of CDK2, specifically phosphorylated CDK2 at critical regulatory residues Y15 and T160, has yet to be defined. To investigate CDK2 isoform levels and the role of activated and inhibited CDK2 throughout prophase I, we have utilised a combination of STA-PUT sorted testis extracts, and immunofluorescence localisation on spread spermatocytes and fixed testis sections.

## Materials and Methods

### Gravitational cell sedimentation (STA-PUT)

STA-put was performed using a modified version^23^ of the original protocol devised by Bellve and colleagues^42^. Eight adult mice (8 weeks) were used in each STA-PUT run. A single cell suspension was made as follows: Testes were dissected and decapsulated into approximately 30 mL of 1 x Krebs buffer (Sigma K3753) supplemented with amino acids (GIBCO 11130-051, Sigma M7145) and glutaMAX (GIBCO 35050-061) in a Petri dish lid. Cell extract was incubated in a shaking water bath at 34°C, 150 rpm in 10 mL of 2 mg/ml Collagenase (Sigma C5138) in 1 x Krebs for 15 minutes. Following three rounds of centrifugation and washing in 20 mL 1 x Krebs, the testis extract was re-suspended in 20 mL of 2.5 mg/ml Trypsin (Sigma T0303), 200 mg/ml DNase (Sigma DN25) in 1 x Krebs and incubated in a shaking water bath at 34°C, 150 rpm for 15 minutes. Extract was then centrifuged and washed three times in 20 mL 1 x Krebs, followed by re-suspension in 20 mL 0.5% BSA (Sigma A7906). Extract was loaded into the STA-PUT apparatus along with 275 mL 4% BSA in 1 x Krebs in the large chamber, 275 mL 2% BSA in 1 x Krebs in the medium chamber and 100 mL 0.5% BSA in 1 x Krebs in the small chamber. Following loading of the BSA into the sedimenting chamber along with the extract, the sample was allowed to sediment for 2 hr. Following sedimentation, cell fractions of 13 mL were collected, centrifuged at 2,000rpm for 5 minutes and washed in 200 µl 1 x PBS. For this STA-PUT run, 41 fractions were collected. For immunofluorescence analysis of cell types 5 µl of each fraction was re-suspended in 5 µl 50mM sucrose and incubated at room temperature for 20 minutes. 30 µl drops of 1% Paraformaldehyde (EMS 19200) were placed on each well of an 8 well slide and the 50 mM sucrose/cell mix were added to each well. Slides were incubated in a humid chamber overnight, followed by drying, washing, and staining against proteins defining stages of prophase I, in our case SYCP3 and γH2AFX. A minimum of 100 cells were scored per fraction. The remaining extract was centrifuged at 2,000 rpm for 5 minutes and re-suspended in 1 x PBS Lysis buffer (1 x PBS, 0.01% NP-40, 5% Glycerol, 150mM NaCl, 1 x Roche cOmplete), sonicated and stored at -20°C for downstream applications.

### Histology and immunofluorescence

Testes were dissected from wildtype adult (8 week old) C57B/6J mice and fixed in 10% neutral buffered formalin for 8 hrs at room temperature. Fixed testes were then washed 4 x in 70% ethanol, embedded in paraffin and sectioned onto glass slides at a thickness of 5 µM. Slides were rehydrated in safeclear followed by decreasing percentages of ethanol (100%, 95%, 80%, 70%, 50%, 30%), washed with deionized water, and then boiled in sodium citrate buffer (10mM Sodium Citrate, 0.05% Tween 20, pH 6.0) for 20 minutes. Following subsequent cooling and washes in PBS, slides were blocked in blocking buffer (1 x PBST (1X PBS + 0.1% Triton), 1% BSA, 3% Goat Serum) for an hour and primary antibody dilutions incubated on the sections for two hours at 37°C. Slides were washed in PBST and incubated with fluorescence-conjugated secondary antibodies for one hour at 37°C, washed with PBST and then mounted using a DAPI/Antifade mix. Slides were imaged on a Zeiss Axiophot microscope with Zen 2.0 software. Each image shown is a single z-slice. Antibody concentrations used can be found in supplemental table 1.

### Meiotic Spermatocyte Spreads

Chromosome spreads were prepared as previously described in ^23^. Briefly, decapsulated testes from wildtype adult (8 week old) C57B/6J mice were incubated in hypotonic extraction buffer (HEB: 30 mM Tris pH7.2, 50 mM sucrose, 17 mM trisodium dehydrate, 5 mM EDTA, 0.5 mM DTT, 0.1 mM PMSF, pH8.2-8.4) for one hour. Small sections of testis tubule were manually dissected into 100 mM sucrose and spread onto 1% Paraformaldehyde, 0.15% Triton X coated slides and incubated in a humid chamber for 2.5 hrs at room temperature. Slides were dried for 30 minutes, washed in 1x PBS, 0.4% Photoflo (Kodak 1464510) and either stored at 80°C or stained. For staining, slides were washed in 1 x PBS, 0.4% Photoflo for 10 minutes, followed by a 10 minute wash in 1x PBS, 0.1% Triton X and finally a 10 minute wash in 10% antibody dilution buffer (3% BSA, 10% Goat Serum, 0.0125% Triton X, 1 x PBS) in 1 x PBS. Primary antibodies were diluted in 100% antibody dilution buffer, placed as a bubble on parafilm within a humid chamber, and the surface of the slide spread on the parafilm allowing the antibody to spread across the surface of the slide. Slides were incubated at 4°C overnight. Slides were washed in 1 x PBS, 0.4% Photoflo for 10 minutes, followed by a 10-minute wash in 1 x PBS, 0.1% Triton X and finally a 10-minute wash in 10% antibody dilution buffer. Secondary antibodies were diluted as the primary antibodies and spread in a similar fashion. Slides were incubated at 37°C for one hour. Slides were washed in 1 x PBS, 0.4% Photoflo for 10 minutes, three times. Finally slides were left to dry, mounted using DAPI/antifade mix and either imaged or stored at 4°C for later imaging. Slides were imaged on a Zeiss Axiophot microscope with Zen 2.0 software. Antibody concentrations used can be found in supplemental table 1.

### Mice

All mice used for this study were handled following federal and institutional guidelines under a protocol approved by the Institutional Animal Care and Use Committee (IACUC) at Cornell University. All mice used in this study were 8-week-old wild type adult males of the C57B/6J strain background, living under 12 hour light/dark cycles.

### Protein extraction

Decapsulated testis extract was re-suspended into 1 x PBS Lysis buffer (1 x PBS, 0.01% NP-40, 5% Glycerol, 150mM NaCl, 1 x Roche cOmplete) and sonicated for 20 s at 22% amplitude in cycles of 0.4 s on and 0.2 s off. Samples were stored at -20°C.

### Quantification of CDK2 and phospho-CDK2 interstitial foci

Interstitial foci were counted manually on 16-bit images of prophase I spreads with all channels merged. Images were adjusted and merged in ImageJ. Fluorescent signal was defined as an interstitial focus if it: colocalised with SYCP3 signals, if it was brighter than the background staining (signal not colocalised with SYCP3), and if it had a “round” appearance/morphology. Due to bright signal of pT160-CDK2 along on sex chromosomes, potential interstitial foci on the pseudoautosomal region of the sex chromosomes were not included in counts and subsequent quantifications. GraphPad Prism version 9.4.1 for Macintosh was used to generate Mean values standard deviation (s.d.) presented in corresponding figures.

### Quantification of CDK2 and phospho-CDK2 telomere staining

Telomere staining was characterised on 16-bit images of prophase I spreads with all channels merged. Images were adjusted and merged in ImageJ. A signal was defined as end staining if it co-localised with the ends of SYCP3 stained synaptonemal complexes (based on staining performed in ^40^) and was brighter than background signal.

### SDS-PAGE, TCE Exposure, Phos-Tag and Western Blotting

Conventional 1.5 mm SDS-PAGE gels were poured as previously described^23^, except for the inclusion of 2,2,2-Trichloroethanol (TCE) (Sigma T54801-100G) to a final concentration of 0.5% within the resolving phase of the SDS-PAGE. For phos-tag gels (FujiFilm AAL-107M), phos-tag was also added to the resolving phase (at the noted concentrations) in addition to MgCl-_2_ at double the concentration of the phos-tag. Protein samples were run as normal, between 100 and 120V. Following sample separation, TCE activation occurred by exposure to UV on a Bio-rad ChemiDoc MP system for 45 to 60 seconds. Gels were imaged at various exposures until maximal signal was detected without saturation of the image. For phos-tag gels, gels were soaked in 50 ml Bio-rad Turbo-Blot transfer buffer supplemented with 10 mM EDTA 3 x for 10 minutes whilst slowly rotating, followed by one wash in 50 ml Bio-rad Turbo-Blot transfer buffer for 10 minutes. Gels were transferred using the Bio-rad Trans Blot Turbo System using the built in transfer program (1.5 mm 10 minute Midi Gel program).

Membranes were blocked in either 5% Non-Fat Dried Milk in 1 x TBST (CDK2 phospho Y15 and T160 antibodies) or 5% BSA in 1 x TBST (all other antibodies) for 30 minutes, then incubated in primary antibodies in 1 x TBST overnight at room temperature. Membranes were washed in 10 ml 1 x TBST for 10 minutes, 3 x and then incubated in secondary antibody in 1 x TBST for 1 hr at room temperature. Membranes were washed in 10 ml 1 x TBST for 10 minutes, 3 x and developed using ECL chemical and imaged on a Bio-rad ChemiDoc MP system. Antibody concentrations used can be found in supplemental table 1.

### TCE and Western Blot Quantification

TCE and western blot images were quantified using Bio-rad ImageLab software. Lanes were detected and total signal recorded from TCE gels. Bands from western blot images were captured and quantified as above. Quantified protein signal was calculated by dividing the western blot signal detected, by the total loaded protein (TCE signal), with the signal plotted.

## Results

### Both CDK2 isoforms are regulated by inhibitory and activatory phosphorylation in the mouse testis

CDK2 in mouse exists as two isoforms (CDK2^33kDa^ and CDK2^39kDa^) due to an alternative splicing event leading to the addition of 48 amino acids between methionine 196 and valine 197 producing CDK2^39kDa^ (figure 1A)^41^. Both isoforms of CDK2 contain multiple residues capable of being phosphorylated (figure 1A), and both can function as active kinases in conjunction with A or E type cyclins. However, the kinase activity of CDK2^39kDa^ *in vitro* is decreased compared with CDK2^33kDa 41^. Given the requirement for CDK2 in prophase I in multiple organisms, we investigated CDK2 protein levels and phosphorylation status in the mouse testis. We used SDS-PAGE with the incorporation of 2,2,2-Trichloroethanol (TCE) which, following UV activation, causes tryptophan-containing proteins to fluoresce^43,44^, providing a readout for total protein loaded on the gel. Equivalent levels of both CDK2 isoforms were detected from whole testis lysate, but the CDK2^33kDa^ isoform showed higher levels of both the inhibitory phosphorylation at tyrosine 15 (pY15) and the activatory phosphorylation at threonine 160 (pT160) compared to the CDK2^39kDa^ isoform (figure 1B, C), suggesting that the CDK2^33kDa^ isoform is more heavily regulated during spermatogenesis. Interestingly both phospho-specific antibodies detected multiple bands associated with CDK2^33kDa^, indicating multiple post-translational modification events occurring on the same protein molecule.

**Figure 1:**
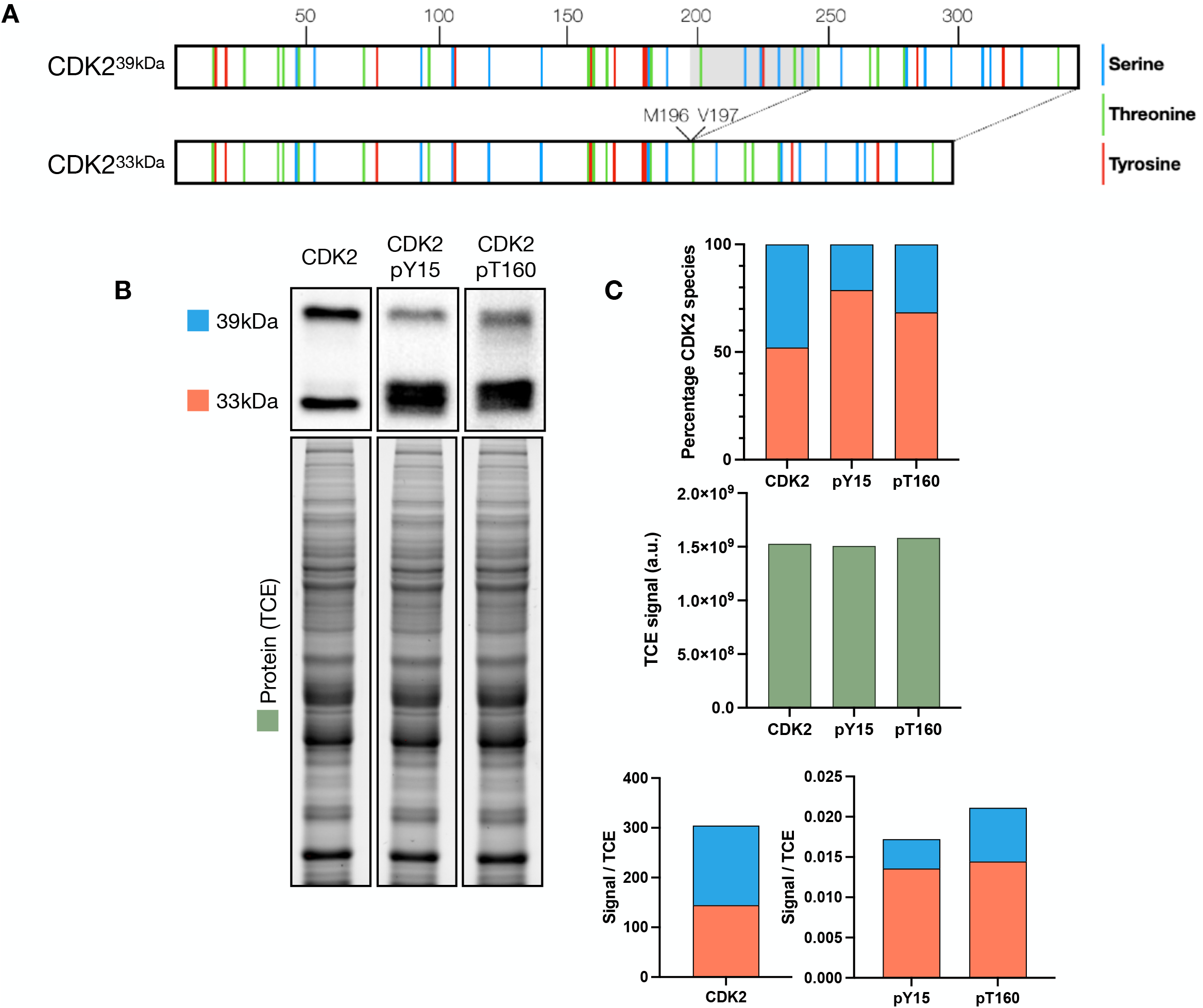
Two CDK2 isoforms are detected in mouse testis and undergo phosphorylation. **A)** Schematic of CDK2 isoforms (CDK2^33kDa^ and CDK2^39kDa^) in mouse with residues capable of being phosphorylated (serine: blue, threonine: green, tyrosine: red) annotated. Position of the alternative splicing region leading to an insertion between M196 and V197 to generate the 39kDa form is displayed with the in-frame region shaded in grey. **B)** Western blots against CDK2, CDK2-phosphoY15 and CDK2-phosphoT160 from a 10% SDS-PAGE. TCE exposure image of western blots indicating total protein loaded. **C)** Quantification of protein detected in B with CDK2 33kDa isoform labelled in red, CDK2 39kDa isoform labelled in blue, and TCE signal detected in green. Top panel depicts the percentage of CDK2 signal from each of the two isoforms, middle panel shows total signal quantified from TCE blots, bottom panel is the signal detected by western blot divided by the TCE protein signal.

### Differential CDK2 isoform protein levels are detected during prophase I

To further investigate meiotic CDK2 function in a stage-specific manner throughout prophase I, we performed gravitational cell sorting (STA-PUT) to isolate specific spermatocyte cell fractions^42,45,46^. STA-PUT separates spermatocytes based on the change in cell size and density that occurs as cells undergo prophase I through loading a single cell suspension from the testes of adult mice to a BSA gradient and allowing the cells to separate over time (figure 2A). Following sedimentation, fractions can be collected and analysed to determine the proportion of each cell type. In this STA-PUT run (figure 2B) we collected 41 fractions and analysed the make-up of each fraction by the morphology of the synaptonemal complex and the sex body (as determined by SYCP3 and γH2AFX staining, respectively). Leptonema enriched cells peaked within fraction 33, zygonema within fraction 28, pachynema within fraction 18, and diplonema within fraction 4. Total protein levels varied across the STA-PUT fractions (as determined by the TCE blot) due to differences in the numbers of cells found within each fraction, reflecting the varying proportion of each cell type in the adult mouse testis.

**Figure 2:**
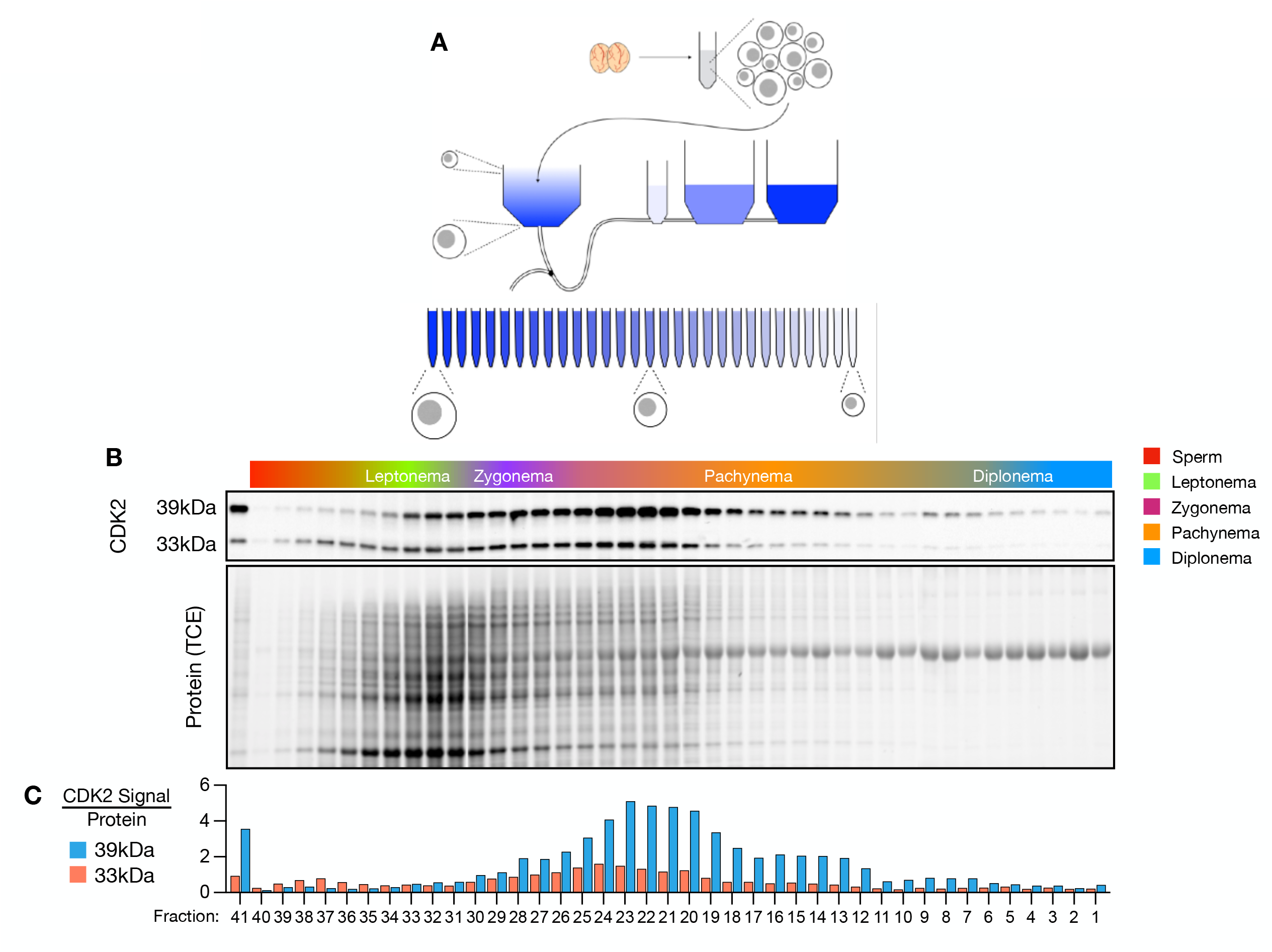
Dynamic CDK2 levels are detected during meiotic prophase I. **A)** Schematic of the STAPUT process. Testes are decapsulated and a single cell suspension generated. Cells are loaded into a collection chamber and different percentages of BSA (blue) flow into the collection chamber. Following gravitational sedimentation cell fractions are collected and analysed. **B)** STAPUT Cell fractions profile following quantification by staining of SYCP3 and γH2AFX with scoring based upon synaptonemal complex and sex body status. CDK2 **w**estern blots and TCE protein gels from STAPUT run are amalgamations of four separate gels, normalised by TCE activation and western blot exposure. **C)** Quantification of protein levels from meiotic STAPUT cell fractions, measured against protein signal from TCE gel exposure (10% SDS-PAGE).

Western blotting across STA-PUT fractions reveals a variation in CDK2 levels throughout prophase I. Interestingly, the two CDK2 isoforms differ in total protein levels, with the CDK2^39kDa^ isoform detected at over three-fold higher than the CDK2^33kDa^ at their peaks, in contradiction to the protein levels observed from whole testis extract where both CDK2 isoforms are detected in almost equal quantities (figure 1B). The dynamics of protein levels between the two isoforms differed in the STA-PUT run, with more CDK2^33kDa^ protein found peaking earlier in prophase I in the zygonema to pachynema fractions, compared with the CDK2^39kDa^ isoform peaking slightly later in the pachynema fractions (figure 2C).

### Phos-tag SDS PAGE allows easier quantification of CDK2 phosphorylation

To determine the amount of phosphorylated versus unphosphorylated CDK2 protein through prophase I, we utilised the phos-tag chemical, which when incorporated in SDS-PAGE gels causes reduced migration of phosphorylated proteins^47,48^. Due to the complexity of handling four gels per condition, potential variability in phos-tag behaviour, TCE activation, and western blotting between membranes, we ran a single gel containing every fourth fraction, to enable the profile of the STA-PUT run to be compared directly. We applied a range of phos-tag concentrations to separate the phosphorylated form due to the complexity of the protein sample. Phos-tag concentrations of 10 µM and 15 µM enabled sufficient band separation to quantify the phosphorylated and unphosphorylated forms and the 15 µM gel was used to quantify protein bands (figure 3A, B). Both CDK2 isoforms showed a similar dynamic of phosphorylation, with a higher proportion of CDK2^33kDa^ observed to be phosphorylated (almost 50% in fraction 29) compared with CDK2^39kDa^ (34%).

**Figure 3:**
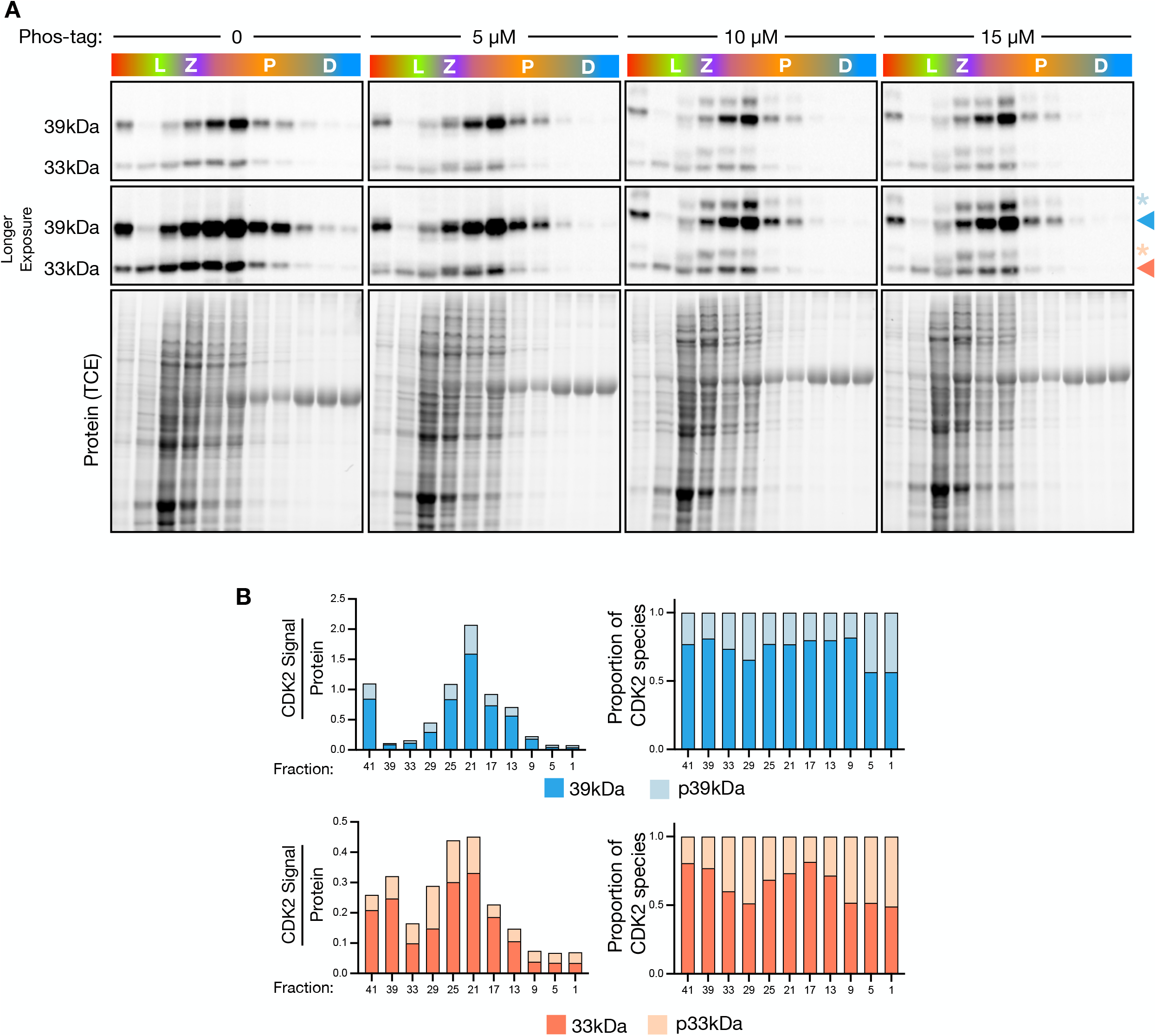
CDK2 phosphorylation can be quantified using phos-tag SDS-PAGE. **A)** CDK2 western blots of STAPUT fractions from 10% phos-tag SDS PAGE gels with concentrations of phos-tag denoted. Middle panel is a longer exposure of the same western blot as the top panel. TCE protein blot from gels used in western blots. **B)** Quantification of CDK2 signal and phosphorylated species from the 15µM phos-tag gel in **A**. CDK2^33kDa^ signal is in red (red arrow) with phosphorylated species in light red/orange (star), CDK2^39kDa^ signal is in blue (arrow) with phosphorylated species in light blue (star).

### STA-PUT fractions reveal dynamic phosphorylation of CDK2 isoforms

Inhibition of CDK2 can occur by phosphorylation at Y15, with activation of CDK2 requiring phosphorylation at T160. To further understand the inhibitory and activatory phosphorylation events on CDK2 during prophase I, we performed western blotting on STA-PUT fractions, probing for pY15 and pT160 (figure 4A). Inhibitory pY15 phosphorylation was detected on both CDK2 isoforms, with more inhibitory phosphorylation detected on CDK2^33kDa^, with the ratios of pY15 across both isoforms remaining similar in most samples (figure 4A, B). To our surprise, minimal activatory pT160 phosphorylation was detected on CDK2^33kDa^ in early STA-PUT fractions, compared with CDK2^39kDa^. pT160 signal of CDK2^33kDa^ only appreciably increases from pachytene-enriched fractions onwards (figure 4A, B).

**Figure 4:**
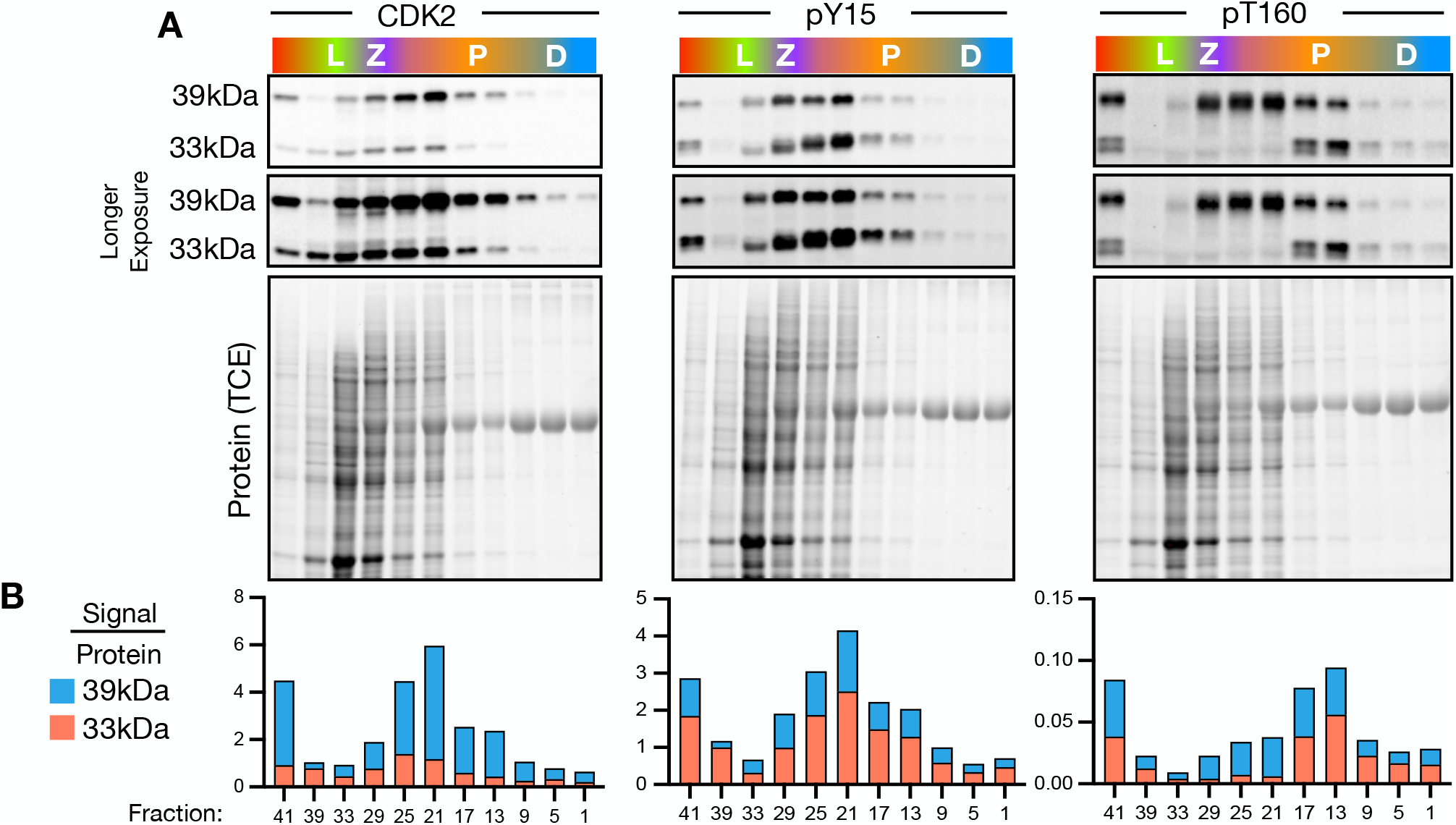
Inhibition and activation of CDK2 isoforms differs throughout prophase I. **A)** CDK2 western blots of STAPUT fractions from 10% SDS PAGE gels. Middle panel is a longer exposure of the same western blot as the top panel. TCE protein blot from gels used in western blots. **B)** Quantification of CDK2 signal from **A**. CDK2^33kDa^ signal is in red, CDK2^39kDa^ signal is in blue.

In addition to phosphorylation at T160, CDK2 activation typically requires partnering with an A or E type Cyclin^26^. Unfortunately, we were unable to source a commercial Cyclin A1 antibody that worked effectively in western blotting. However, western blotting of Cyclin A2 and Cyclin E1 revealed similar dynamics of protein levels during prophase I, with Cyclin E2 observed to peak earlier (Supplemental figure 1). These observations were surprising, given the Cyclin A2 reported functions earlier in prophase I^49^. CDK2 inhibitory phosphorylation at Y15 takes place via the WEE1 kinase^50,51^. WEE1 protein levels on STA-PUT fractions were highest in early fractions and decreased as prophase I progresses (Supplemental figure 1), consistent with a role in inhibiting cell cycle progression due to unrepaired meiotic DSBs and until correct crossover numbers were formed, as we previously reported^23^.

### Activatory T160 phosphorylation foci are detected in pachytene spermatocytes

To further investigate the functions of the inhibitory and activatory phosphorylation events during prophase I, we stained for pan-CDK2, pY15 and pT160 on spread spermatocytes. Consistent with previously published work, we observe CDK2 protein localising to telomeres at zygonema, persisting through to diplonema (figure 5A)^25^. In pachynema, CDK2 localises as interstitial foci along the synaptonemal complex, the vast majority of which co-localise with MLH3, supporting the idea that these are crossover specific foci (figure 5B). Inhibitory Y15 phosphorylation localises to telomeres at zygonema, but by pachynema pY15 signal was only detected along the sex chromosomes and is not found either at the telomere or the interstitial sites on the autosomes. We did not detect activatory T160 phosphorylation at telomeres of zygonema spermatocytes, but at pachynema we detect T160 signal co-localising with SYCP3 along the length of the sex chromosomes, and in the interstitial regions along the autosomes (figure 5A, enlarged panel). The interstitial T160 foci occur at a similar frequency to MLH3, with approximately 70% co-localising with MLH3 (figure 5A enlarged panel, B).

**Figure 5:**
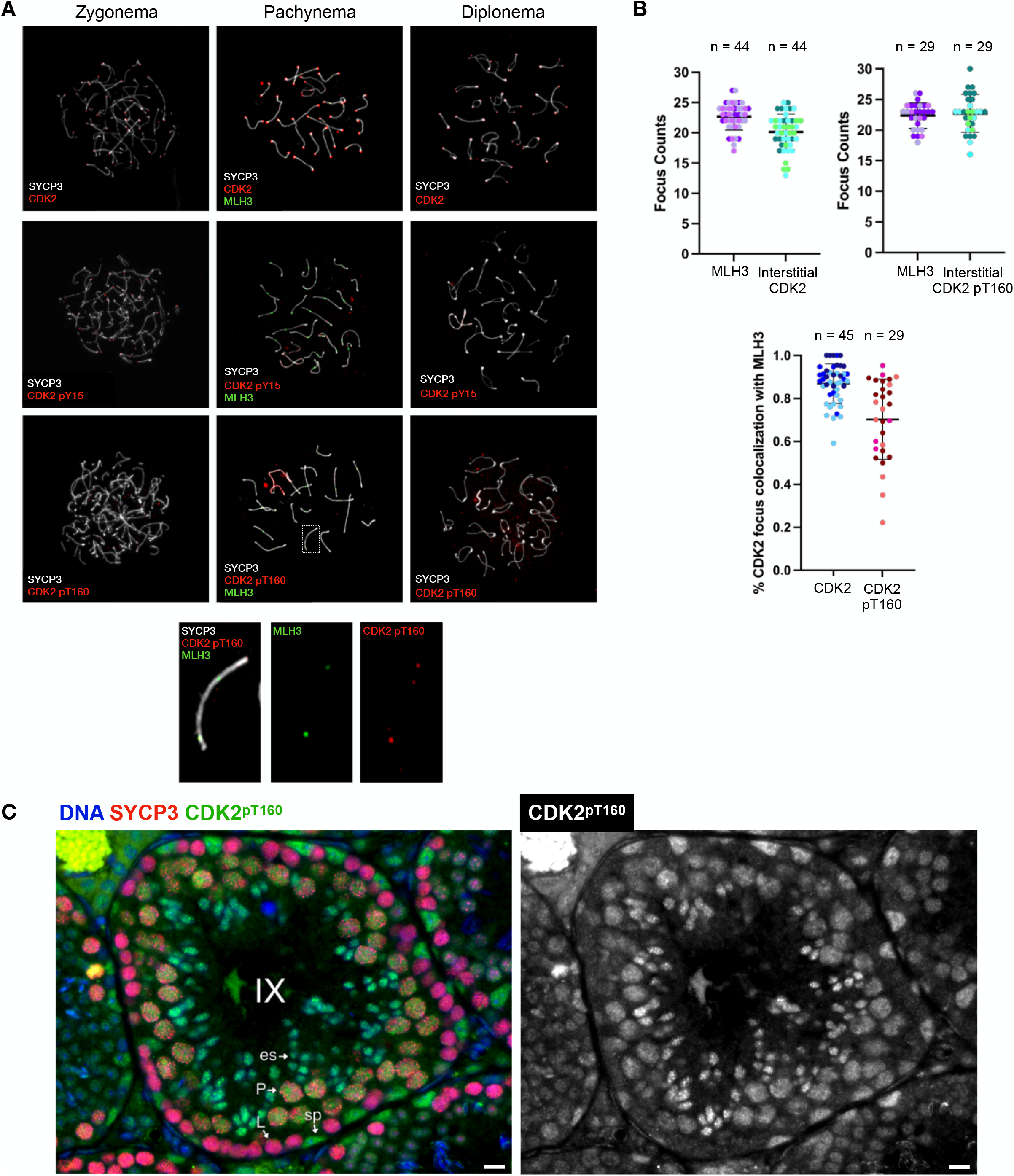
CDK2-pT160 signal localises to crossover sites and is increased in pachytene spermatocytes. **A)** Representative images of spread spermatocytes from wild type C57BL/6J adult males stained for SYCP3 (white) CDK2/pY15/pT160 (red) and MLH3 in zygonema, pachynema and diplonema. Bottom panel are enlargements of pachytene spermatocyte staines with SYPC3, CDK pT160 and MLH3 and separate channels **B)** Quantification of stained spermatocyte spreads. Each dot indicates a cell, with each colour indicating a mouse. Mean plotted with standard deviation **C)** C57BL/6J testes stained for DNA (blue), SYCP3 (red), and CDK2 pT160 (green). A stage IX tubule is show. CDK2 pT160 was not present in leptotene spermatocytes (L) but is enriched in pachytene spermatocytes (P) and elongating spermatids (es). CDK2 pT160 was also found in spermatogonia (sp). Scale bar: 10µm.

Next, we stained fixed testis sections to gain a perspective of activatory T160 phosphorylation through spermatogenesis. pT160 signal was detected in spermatogonia, pachytene spermatocytes and elongating spermatids but absent from leptotene spermatocytes (figure 5C), suggesting that CDK2 pT160 is absent as primary spermatocytes begin meiosis, with CDK2 activation occurring in pachynema.

## Discussion

The current study was aimed at elucidating critical dynamic profiles for the two CDK2 isoforms through meiotic prophase I in mouse spermatocytes. In addition, we set out to elucidate the relative levels of inhibitory and activatory post-translational modifications on CDK2, focusing on pY15 and pT160, respectively. Using a combination of western blotting, localisation of protein foci on meiotic spread chromosome preparations, and immunofluorescence on paraffin-embedded testis sections, this work has uncovered differential isoform expression and tightly controlled temporal and spatial inhibition/activation of CDK2 during prophase I, specifically with regards to the two CDK2 isoforms (figure 6). Specifically, we observe peak levels of CDK2^33kDa^ between zygonema and pachynema, with CDK2^39kDa^ peaking slightly later in earlier pachynema and at higher levels. Both isoforms of CDK2 show similar kinetics of inhibitory pY15 phosphorylation, but surprisingly whilst the activatory pT160 phosphorylation of CDK2^39kDa^ is similar to that of the pY15 phosphorylation, CDK2^33kDa^ phosphorylation only occurs in mid pachynema, coinciding with the timing of crossover formation (figure 6).

**Figure 6:**
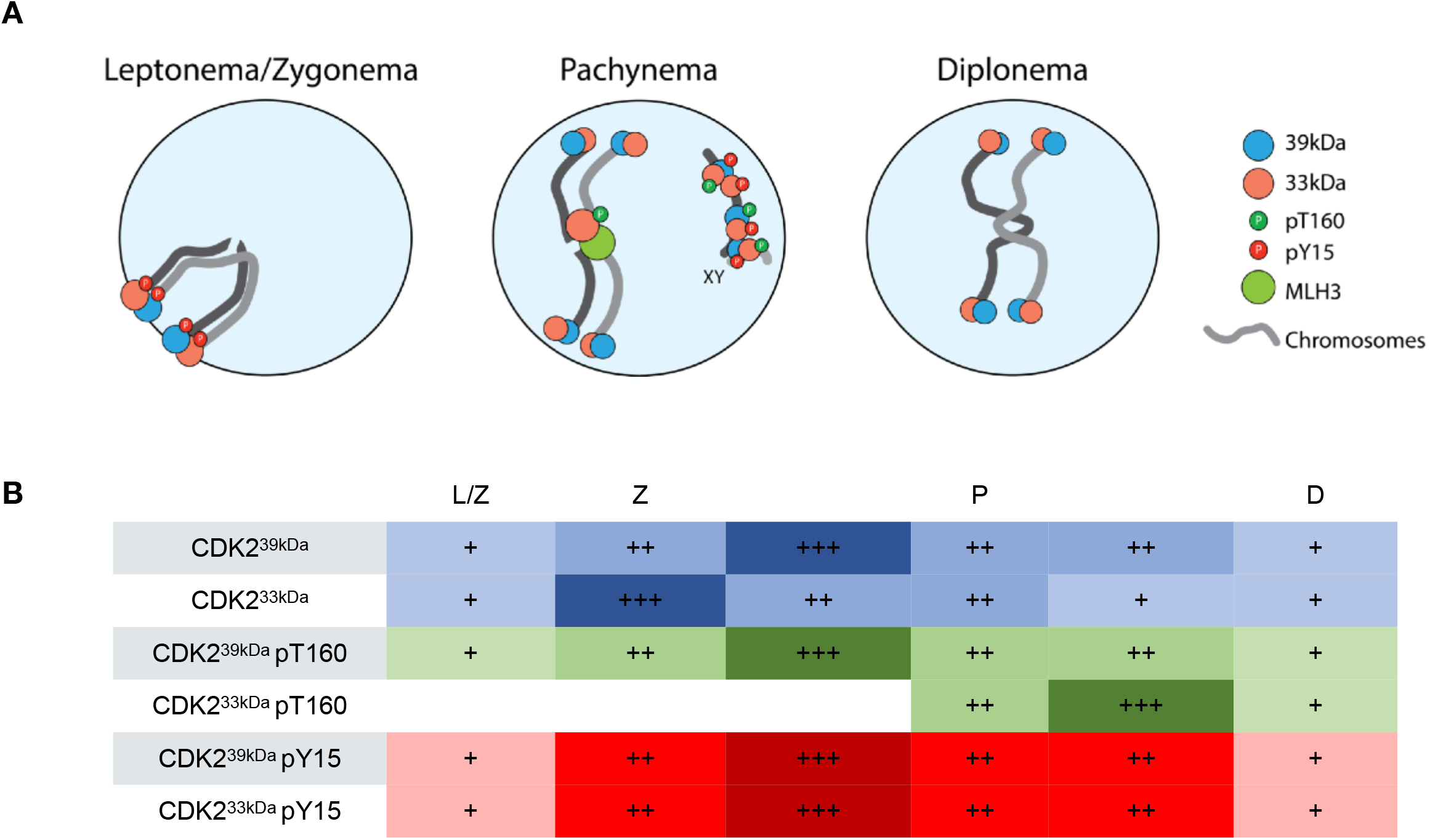
Summary of CDK2 isoform and phosphorylation levels, and localisation through prophase I. **A)** Localisation of CDK2 through prophase I. Inhibitory pY15 CDK2 and pan-CDK2 localises to telomeres during leptonema, with CDK2 and activated pT160 CDK2localises to interstitial sites, colocalising with MLH3 at pachynema. Inhibited and activated CDK2 localises along the sex chromosomes. By diplonema, only CDK2 signal is detected at the telomeres. **B)** Summary of CDK2 protein levels. Levels are expressed as plus signs, based upon signal detected by western blotting. Protein levels are expressed in blue, activatory T160 phosphorylation in green and inhibitory Y15 phosphorylation in red. L/Z: Leptonema/Zygonema, Z: Zygonema, P: Pachynema, D: Diplonema.

Our investigations into CDK2 from whole testis lysate revealed multiple bands when using the phosphorylation specific antibodies. Given the low signal detected for both phosphorylated species relative to total CDK2 (figure 1B, C bottom panel), it is likely these bands are of too low abundance to detect relative to the unmodified signal when using a pan-CDK2 antibody. Interestingly, it is the CDK2^33kDa^ isoform that shows multiple post-translational modification events and therefore further regulation. Together with the kinetics observed of the T160 phosphorylation, it supports the idea that the important function isoform of CDK2 during prophase I is the CDK2^33kDa^ isoform. The inhibition and activation of CDK2 phosphorylation at Y15 and T160, respectively, have previously been described^35^ and we observe these phosphorylation events occurring dynamically during prophase I. In addition to Y15 and T160, other phosphorylation events have been described on CDK2, notably T14, as a similar inhibitory mechanism^35^, and phosphorylation on T158 and Y159 based upon bioinformatic predictions. These additionally described residues capable of being phosphorylated, may account for the additional bands observed on the CDK2^33kDa^ isoform. Interestingly in our analysis of CDK2 from whole testis lysate and from STA-PUT isolated fractions, the abundance of the CDK2 isoforms changed from approximately equal amount of each in whole testis (figure 1B, C), to CDK2^39kDa^ being dominant in STA-PUT fractions (figure 2). Whilst we do not understand why this is the case, whole testis lysate contains protein from the all cells of the testis, including blood and Sertoli cells which are likely filtered out during the STA-PUT cell preparation. This intriguing difference, however, may have implications when investigating CDK2 function using whole testis lysate, as differentially expressed isoforms at specific stages of germ cell development versus testis vasculature and support cells may confound experimental outcomes.

Our analysis of CDK2 phosphorylation throughout prophase I revealed a surprising temporal regulation of the two CDK2 isoforms (summarised in figure 6). Western blotting revealed the levels and kinetics of two CDK2 isoforms differed (figure 2B, C, 6B), most notably in the activatory phosphorylation of CDK2^33kDa^ (figure 4A, B, 6B). Specifically, both spread spermatocyte and testis section staining of pT160 revealed the onset of signal at pachynema, localising to crossover sites (figure 5, 6A). Thus, there is a stage-specific phosphorylation event at pachynema that coincides with the designation of the final tally of Class I crossovers. This important temporal regulation of CDK2 provides potential insights into the signalling cascades that occur following the formation of crossover structures. Importantly, whilst CAK has been previously described as activating CDK2 by phosphorylation on T160^36^, it has yet to be determined if this is kinase responsible for phosphorylation during pachynema and, if so, how indeed this kinase is also regulated. Not only has the T160 staining provided us with important insight into CDK2 regulation in the context of class I crossovers, but it has also provided a unique tool that can be used to accurately stage different prophase I stages. Antibodies against the unique region of CDK2^39kDa^ would enable us to investigate if the important active form of CDK2 during prophase I is the CDK2^33kDa^ isoform. Given that the *Cdk2*^*-/-*^ and kinase dead mice arrest in zygonema^31-33^, the activatory T160 phosphorylation of CDK2^33kDa^ at pachynema would correlate with this being an important stage of germ cell development, indicating a developmental transition between zygonema and pachynema dependent upon CDK2 activity. Given the localisation of CDK2 pT160 to crossover sites at pachynema, and crossover formation being the defining developmental process occurring during pachynema, this would provide strong evidence that CDK2 is playing an important role at recombination sites. Identification of the targets of CDK2 would elucidate this function and provide important mechanistic regulation into prophase I events.

## Data Availability

The data that support the findings of this study are contained within the article and the supporting information. All source data generated for this study are available from the corresponding author (Stephen Gray) upon reasonable request.

