## Supplemental Figures for "Dynamic regulatory phosphorylation of mouse CDK2 occurs during meiotic prophase I"

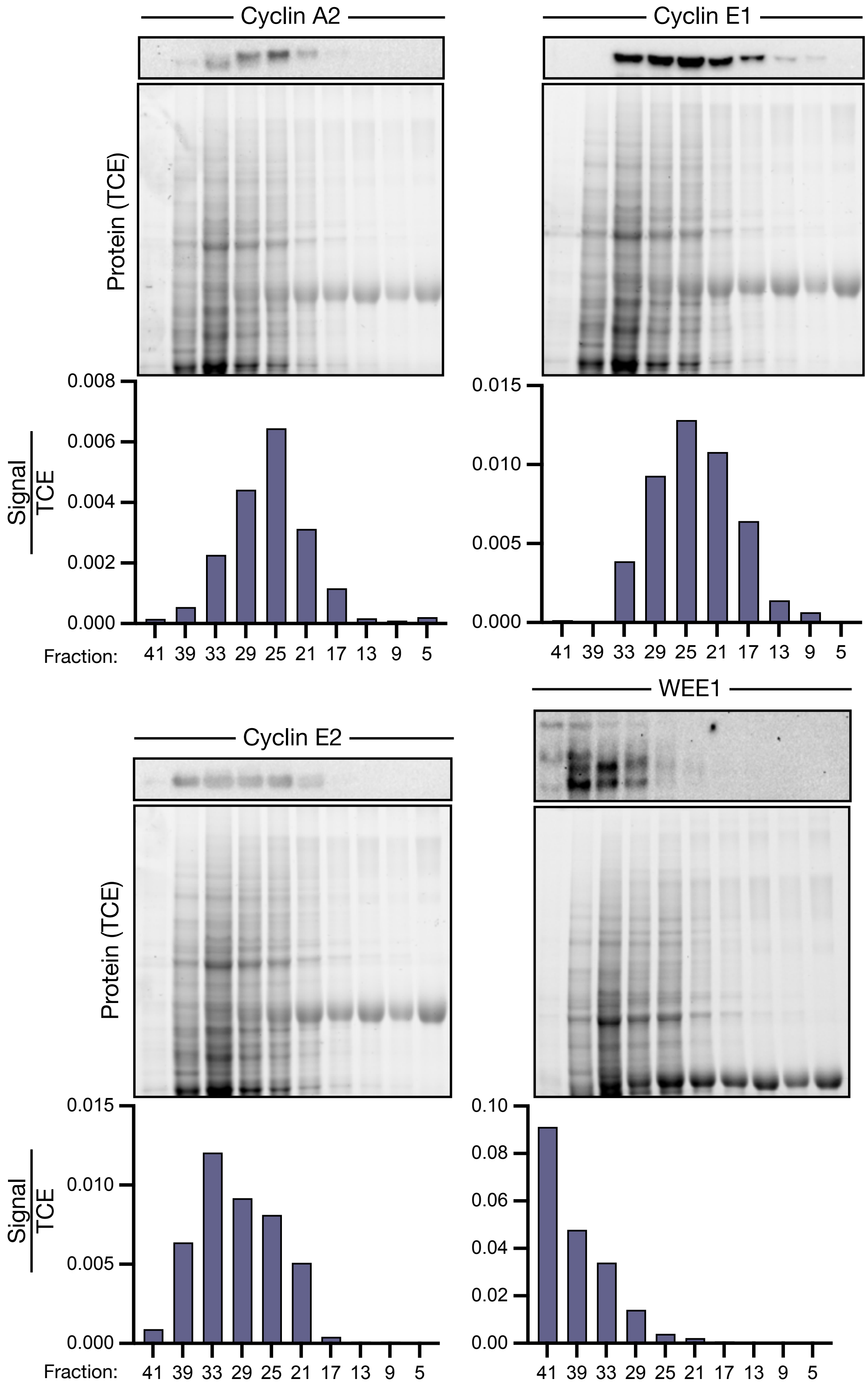


**Supplemental Figure 1: Protein levels of CDK2 interacting Cyclins and the CDK2 inhibitor WEE1.** Western blots of STAPUT fractions with TCE protein blot from gels used in western blots. Quantification of signal presented.

| Antibody | Organism | Supplier | Product Code |
|---|---|---|---|
| CDK2 | Rabbit | Santa-Cruz | sc-163 |
| CDK2 | Mouse | Santa-Cruz | sc-6248 |
| CDK2 | Rabbit | Protein Tech | 10122-1-AP |
| CDK2 pY15 | Rabbit | Abcam | ab76146 |
| CDK2 pT160 | Rabbit | Abcam | ab194868 |
| SYCP3 | Mouse | Abcam | ab97672 |
| SYCP3 | Rabbit | Custom home made | - |
| MLH3 | Guinea Pig | Custom home made | - |
| Cyclin A2 | Rabbit | Protein Tech | 18202-1-AP |
| Cyclin E1 | Rabbit | Protein Tech | 11554-1-AP |
| Cyclin E2 | Rabbit | Protein Tech | 11935-1-AP |
| WEE1 | Rabbit | Cell Signaling | 4936S |

**Supplemental Table 1: Antibodies used in this study**
